# IL-17A promotes epithelial cell IL-33 production during nematode lung migration

**DOI:** 10.1101/2022.11.03.515050

**Authors:** Jesuthas Ajendra, Stella Pearson, James E. Parkinson, Brian H.K. Chan, Henry J. McSorley, Tara E. Sutherland, Judith E. Allen

## Abstract

The early migratory phase of pulmonary helminth infections is characterized by tissue injury leading to the release of the alarmin IL-33 and subsequent induction of type 2 immune responses. We recently described a role for IL-17A, through regulation of IFNγ, as an important inducer of type 2 responses during infection with the lung-migrating rodent nematode *Nippostrongylus brasiliensis*. Here, we aimed to investigate the interaction between IL-17A and IL-33 during the early lung migratory stages of *N. brasiliensis* infection. In this brief report, we demonstrate that deficiency of IL-17A leads to impaired IL-33 expression and secretion early in infection, independent of IL-17A suppression of IFNγ. Impaired IL-33 production was evident in lung epithelial cells, but not innate immune cells. Therefore, our results demonstrate that IL-17A can drive IL-33 during helminth infection, highlighting an additional mechanism through which IL-17A can regulate pulmonary type 2 immunity.

## Introduction

Interleukin-33 (IL-33) belongs to the IL-1 cytokine family and plays a key role in innate and adaptive immunity. IL-33 signals via the ST2 receptor, which is expressed on many different cell types including eosinophils, group 2 innate lymphoid cells (ILC2) and Th2 cells^1^. The expression of ST2 on cell types closely associated with type 2 immunity and evidence that IL-33 is a potent inducer of type 2 responses makes the IL-33/ST2 axis a therapeutic target in type 2 mediated diseases. However, IL-33 is implicated more broadly in the maintenance of tissue homeostasis, and has roles in protection against microbial infection and regulatory T cell (Treg) expansion^2^. In contrast to most other cytokines, IL-33 is usually released during cellular necrosis, underpinning its role as an “alarmin” cytokine^3^. However, studies have also demonstrated IL-33 release by cells of the airway epithelium including alveolar epithelial type II (ATII) cells after infection with *Strongyloides venezuelensis* and administration of *Alternaria alternata*, respectively^4,5^.

The proinflammatory cytokine IL-17A is typically associated with host protection against fungal and bacterial infections^6^. However, using a model of infection with the lung-migrating nematode *Nippostrongylus brasiliensis* (*Nb*), we recently demonstrated that IL-17A is also necessary to mount a pulmonary type 2 response^7,8^. In this helminth setting, IL-17A from innate γδ T cells exerts a suppressive effect on IFNγ release by multiple cell types early during infection^7^. This inhibition of IFNγ allows development of the adaptive pulmonary Th2 response. However, existing data on the interaction between IL-17A and IL-33^9–12^ suggests the possibility that innate IL-17A during *Nb* infection may function beyond its ability to suppress IFNγ. In a neonatal mouse model of influenza, infection-induced IL-17A was associated with increased IL-33 production by lung epithelial cells, subsequently generating a local type 2 immune response^9^. Similarly, mice lacking IL-17A exhibit decreased IL-33 in visceral adipose tissue, leading to reduced Treg expansion and failure to regulate thermogenesis^10^. In both these studies, as in lung *Nb* infection, the source of IL-17A is γδ T cells^7,8^. IL-17A-induced IL-33 also promotes type 2 immunity during atopic dermatitis with the IL-17A source being ILC3s^11^. Of note, during pulmonary *Aspergillus fumigatus* infection, IL-33 negatively regulates IL-17A production^12^, demonstrating cross-regulation between these two cytokines. During *Nb* infection, we have shown IL-17A to be a driver of type 2 immunity^7,8^, but Hung et al. have demonstrated a pivotal role for IL-33 in the type 2 response in *Nb* infection^13^. Given the existing literature on the interactions between IL-17A and IL-33 described above, we felt it essential to investigate the relationship of these two cytokines during the lung-migrating phase of *Nb* infection.

In this brief report we describe IL-17A stimulation of IL-33 release by the lung epithelium during the early phase of *Nb* infection. However, this effect was independent of the IFNγ-suppressing function of IL-17A. Furthermore, IL-17A-induced neutrophilia and the associated lung damage was not the driver for IL-33 production. Together, our data demonstrate that IL-17A acts as an upstream regulator of type 2 immune responses in the lung through two distinct pathways; IFNγ-suppression and IL-33-secretion.

## Material & Methods

### Mice

C57BL/6 J mice were obtained from Charles River. C57BL/6 *Il17a*^Cre^*Rosa26*^eYFP^ mice were originally provided by Dr Brigitta Stockinger^14^. C57BL/6 *Il17a*^Cre^*Rosa26*^eYFP^ homozygote mice are IL-17A-deficient and described here as *Il17a*-KO. Male and female mice were age- and sex-matched and housed in individually ventilated cages. Experimental mice were not randomized in cages, but each cage was randomly assigned to a treatment group. Mice were culled by asphyxiation in a rising concentration of CO_2_. Experiments were performed in accordance with the United Kingdom Animals (Scientific Procedures) Act of 1986 (project license number 70/8547).

### *N. brasiliensis* infection

*Nb* was maintained by serial passage through Sprague-Dawley rats, as described^15^. Third-stage larvae (L3) were washed ten times with PBS (Dulbecco’s PBS, Sigma) before infection. On day 0, mice were infected subcutaneously with 250 larvae (L3). At various time points mice were euthanized and lungs were taken for further analysis.

### Flow cytometry

Single-cell suspensions of the lung were prepared for flow cytometry as previously described^7^. For intracellular staining of IL-33 (clone: 396118, Invitrogen) by epithelial cell adhesion molecule (EpCAM) positive cells, a previously described protocol was used^16^. Cells were stimulated for 4 h at 37 °C with cell stimulation cocktail containing protein transport inhibitor (eBioscience), then stained with live/dead (Thermo Fisher Scientific). After surface antibody staining, cells were fixed for o/n at 4° C using ICC Fixation buffer (eBioscience), then incubated for 20min at RT in permeabilization buffer (eBioscience). Intracellular staining was performed for IL-33 for 30min at RT. Samples were acquired with an LSR Fortessa II flow cytometer and data analysed using FlowJo software.

### Histology and Immunofluorescence

For histology, lungs were inflated with installation of 10% neutral-buffered formalin (NBF; Sigma) and the left lobe of the lung was isolated following *Nb* infection and submersion fixed in NBF. Whole left lung lobes were processed using a tissue processor (Leica ASP300S) and embedded in paraffin. Paraffin blocks were then sectioned to 5 μm using a microtome (Leica RM2235). For immunostaining, lung sections were deparaffinised, rehydrated and heat-mediated antigen retrieval using 10mM sodium citrate buffer (pH 6.0) was performed. To block non-specific protein-binding, 10% donkey serum was used. Sections were stained using an IL-33 antibody (ab118503, Abcam) 1:1000 diluted in PBS followed by a secondary fluorochrome anti-rabbit NorthernLights NL637-conjugated antibody (NL005; R&D systems, 1:200). Sections were then mounted with Fluormount G containing DAPI (Southern Biotech). Staining was visualised on an EVOS^™^ FL Imaging System (ThermoFisher Scientific) and fluorescence intensity was calculated with ImageJ software (version 1.44) as described before^17^. For all images background was subtracted, then images were brightened to a minimum threshold. IL-33-KO mice, which were kindly provided by Dr. John Grainger (Lydia Becker Institute, Manchester, UK) were used to validate specificity of antibody staining.

### Neutrophil depletion

Neutrophil depletion was performed as described before^7^. In short, mice were injected intraperitoneally with 500 μg of neutrophil depleting antibody (clone 1A8, BioXcell) days –1 and 1 post infection. Control mice were treated with a corresponding isotype control (IgG2a).

### IL-33 inhibition

IL-33 was suppressed using *Heligmosomoides polygyrus* alarmin release inhibitor (HpARI). HpARI was generated as previously described^18^. HpARI (10μg) was administered intranasally in 30 μl of PBS on day 0 and day 1 post infection with *Nb*. Controls were treated with 30 μl PBS intranasally.

### mRNA quantification

RNA extraction and quantitative rtPCR was performed as previously described^7^. The expression of *Il33* mRNA was normalized to that of the housekeeping gene *Actb* (β-actin).

### IL-33 ELISA

Bronchoalveolar lavage (BAL) was performed using 10% FBS in PBS. Collected BAL fluid was centrifuged at 400g for 5min to pellet the cells, and supernatants collected. IL-33 concentrations were measured by ELISA (IL-33 duoset, R&D systems) and detected using horseradish peroxidase-conjugated streptavidin and TMB substrate (Biolegend) and the reaction stopped with H2SO4. Absorbance was measured at 450nm using a VersaMax microplate reader (Molecular Devices).

### Statistics

Prism 7.0 (version 7.0c, GraphPad Software) was used for statistical analysis. Differences between experimental groups were assessed by Kruskal-Wallis test for nonparametric data, followed by Dunn’s multiple comparisons test. For gene expression data, values were log2 transformed to achieve normal distribution. Comparisons with a P value < 0.05 were considered to be statistically significant. Data are represented as mean ± sem.

## Results & Discussion

### Lack of IL-17A leads to impaired early IL-33 production during *Nb* infection

IL-33 is a key alarmin strongly associated with type 2 immunity^19^. Both IL-33 and its receptor ST2 have been shown to be implemented in the defense against different nematode infections including *Nb*^13^, *Litomosoides sigmodontis*^20^ and *Strongyloides ratti*^3^. We recently described IL-17A as an important initiator of type 2 responses in the lung by downregulating early IFNγ production^7^. To investigate the relationship between IL-17A and IL-33 during the early phase of infection, both *Il17a*-KO and WT mice were infected with 250 L3 *Nb* larvae and the pulmonary immune response assessed at d1 and d2pi. Using flow cytometry, we found that infection driven IL-33 was produced by EpCAM^+^CD45^−^ lung epithelial cells (Fig. 1A). A comparison between WT and *Il17a*-KO mice demonstrated a significant reduction in IL-33 production by EpCAM^+^CD45^−^ cells in *Il17a*-KO mice compared to WT controls ((Fig. 1B). IL-33 expression was detected in the lung epithelium early during infection at 16h (Fig. 1C), possibly correlating with the time point *Nb* larvae enter the lung. Using ELISA, we determined that by 24h post infection WT mice secreted significantly higher amounts of IL-33 into the bronchoalveolar lavage fluid (BAL) compared to infected *Il17a*-KO mice (Fig. 1D). Taken together, these data reveal an IL-33 regulating function of IL-17A during the early stages of *Nb* infection.

**Figure 1:**
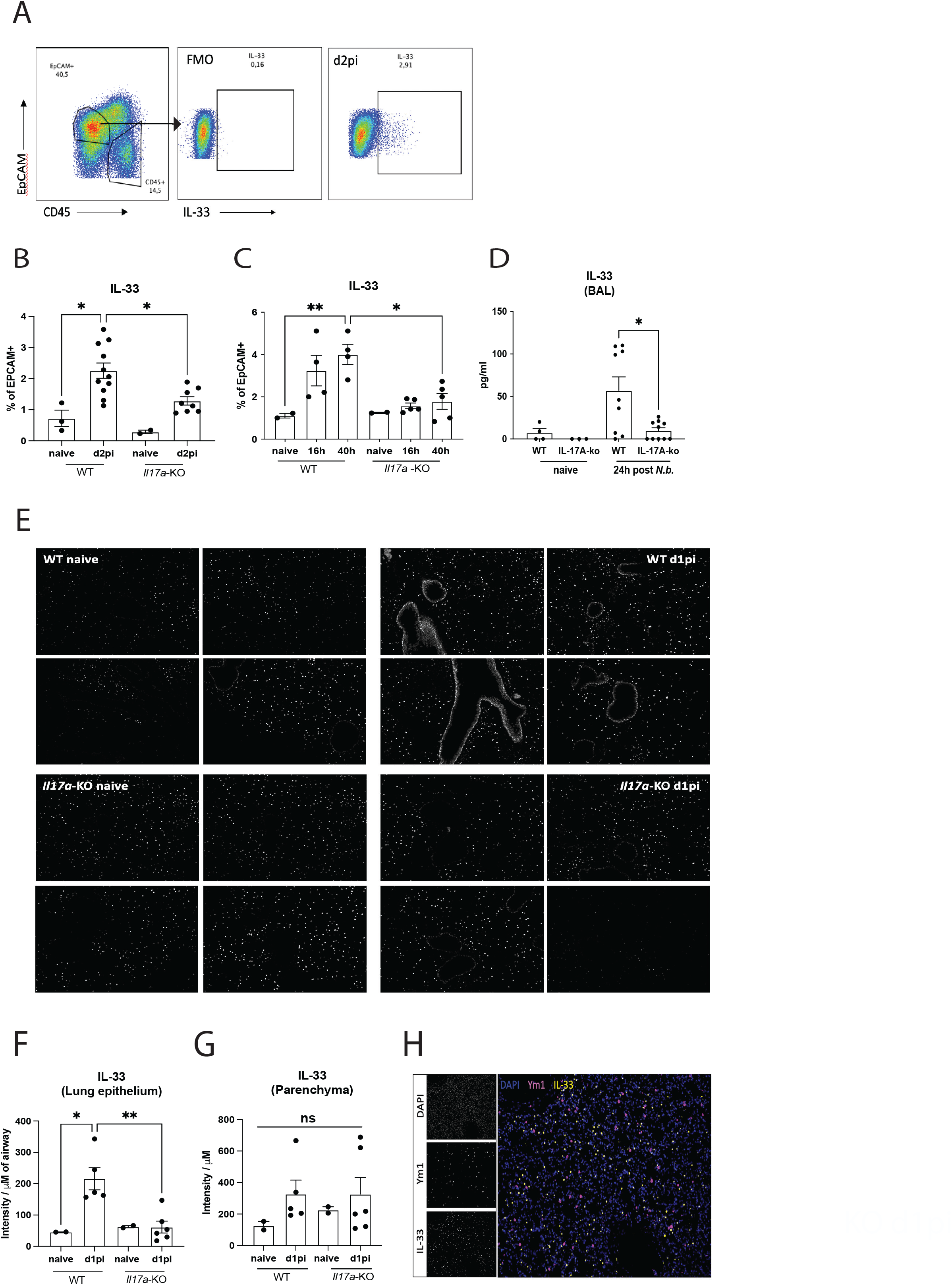
Mice were infected with 250L3 larvae *Nb* and IL-33 production was assessed using different techniques. Representative flow cytometry gating strategy showing intracellular IL-33 for infected mice. Whole lung cells were gated on single and live cells, then for Epcam+CD45-lung epithelial cells (A). Frequency of intracellular IL-33 in Epcam+CD45- lung epithelial cells in WT and *Il17a*-KO mice d2pi (B) and 16h and 40h post *Nb* infection (C). IL-33 in BAL fluid of naïve and mice 24h post infection (D). Microscopy images of immunofluorescent staining for IL-33 (white) in lung sections of WT and *Il17a*- KO mice 1d post *Nb* infection and naïve controls (E). Quantification for IL-33 positive areas for lung epithelium (F) and lung parenchyma (G). Antibody positive staining area was quantified for the intensity normalized to background staining. Immunofluorescent staining for Ym1 (pink) and IL-33 (yellow) in lung sections of WT mice to confirm co-staining (H). Data are expressed as mean ± s.e.m. and are representative for 2 individual experiments and were analysed by ANOVA with Tukey’s multiple comparison test. \**P* < 0.05, \*\**P* < 0.01.

To further identify localization of IL-33 production during *Nb* infection, we examined immunofluorescence staining in lung sections from d1 infected WT and *Il17a*-KO mice. IL-33 expression was detected in lungs of both, naïve WT and *Il17a*-KO mice (Fig. 1E). However, consistent with the flow cytometry data the bronchial epithelial cell staining for IL-33 was less intense in *Nb* infected *Il17a*-KO mice, relative to the infected WT mice (Fig. 1E). In accordance with the observations of Hung et al.^13^, quantification of the IL-33 positive areas of the lung airways revealed a significant increase in IL-33 intensity between naïve and d1 infected WT mice (Fig. 1F). However, we observed significantly lower IL-33 intensity in the airways of *Il17a*-KO mice (Fig. 1F). In fact, *Il17a*-KO mice failed to upregulate IL-33 in the bronchial epithelium after *Nb* infection. In contrast to the bronchial epithelium, IL-33 expression by the cells in the lung parenchyma, did not differ between genotypes at the same time point. IL-33 intensity in the lung parenchyma was quantified, but we did not observe any statistical differences between all tested groups (Fig. 1G). Due to previous reports^16,22^, we investigated the possibility that myeloid cells were the source of IL-33 expression in the lung parenchyma by co-staining lung sections for both IL-33 and Ym1. Ym1 is a chitinase-like protein known to induce IL-17A and thereby promote pulmonary type 2 responses^8^. In the lung, Ym1 is mainly expressed by myeloid cells including macrophages and neutrophils^23,24^. Lung sections stained for both Ym1 (pink) and IL-33 (yellow) revealed no co-staining, suggesting that the source of IL-33 in the parenchyma was not myeloid cells (Fig. 1H). The lung epithelium is a major barrier surface and is known to respond to IL-17A by producing antimicrobial proteins or neutrophil chemoattractants^25^. As such, lung-epithelial specific deletion of IL-17RA results in impaired clearance of *Klebsiella pneumoniae* infection^26^. Our finding that during early *Nb* infection, the presence of IL-17A leads to increased airway epithelial derived-IL-33, along with other recent studies^9^ suggest that IL-17A regulation of lung epithelium extends beyond the induction of proinflammatory and anti-microbial factors, and involves regulation of type 2-associated molecules.

The molecule Ym1 is responsible for early IL-17A production during *Nb* infection^8^, and thus an early type 2 ‘trigger’ may be needed for IL-17A to promote type 2 responses. The early Ym1 needed to induce IL-17A and promote type 2 immunity, is mostly IL-4Rα independent^23^ but as infection progresses Ym1 production becomes increasingly IL-4Rα-dependent, and functions to repair lung damage, in part through the induction of RELMα by lung epithelial cells^23^. At this stage, when the type 2 immune response is fully established, the role of IL-17A switches to suppress rather than enhance the type 2 immune response. Whether IL-17A regulation of IL-33 also differs during the early versus later adaptive stages of infection is unknown, but Hung et al have demonstrated that the context and cell source of IL-33 impacts its role in anti-helminth immunity^22^. Thus, it is important to understand the specific consequence of epithelial cell-derived IL-33. A possible mechanism by which IL-17A induces IL-33 is suggested by work on human epidermal keratinocytes, where IL-17A causes phosphorylation of EGFR, ERK, p38 and STAT1, which is necessary for the induction of IL-33^27^.

### Suppressing IL-33 during *Nb* infection does not increase IFNγ production

We have reported that during *Nb* infection, early IL-17A is responsible for the suppression of IFNγ from all cellular sources^7^. Here we observed that in the absence of IL-17A IL-33 expression in the lung epithelium and IL-33 secretion into the BAL fluid is reduced. We therefore asked whether the ability of IL-17A to limit IFNγ production is acting indirectly through promotion of IL-33. To answer this question we used a known suppressor of IL-33, the *H.polygyrus*-derived protein HpARI, previously shown to inhibit type 2 immune response during *Nb* infection^18^. HpARI directly binds to IL-33 and nuclear DNA, blocking the interaction of IL-33 with the ST2 receptor^18^. Furthermore, HpARI prevents release of IL-33 from necrotic cells^18^. HpARI was administered intranasally at d0 and d1 of *Nb* infection and IFNγ production was investigated d2pi (Fig. 2A). Consistent with our previous findings^7^, at d2pi there was a significant decrease in the proportion of CD8+ T cells (Fig. 2B), γδ T cells, and NK cells producing IFNγ (Fig. 2B-E). Notably, blocking IL-33 with HpARI in *Nb* infected mice did not reverse the suppression of IFNγ (Fig. 2 B-E). In contrast and as expected, *Il17a*-KO mice exhibited significantly increased IFNγ production compared to WT controls at d2pi with *Nb*. To validate that HpARI treatment was successfully blocking IL-33, we analyzed the numbers of ILC2s in the lung and observed a significant reduction in IL-5+ILC2s in the lung at d2pi (Fig. 2F) further confirming the requirement for IL-33 in type 2 mediated immunity during *Nb* infection^13^. Additionally, IL-33 protein levels in lung homogenates were significantly reduced after HpARI treatment (Fig. 2G). In summary, the evidence does not support a role for IL-33 in the suppression of IFNγ during the early stages of pulmonary nematode infection. Because blocking IFNγ rescues type 2 immunity in *Il17a*-KO mice^7^ the suppression of IFNγ by IL-17A is sufficient to initiate type 2 responses, without the need to enhance IL-33 expression. The data therefore suggests that IL-17A induction of type 2 responses involves two fully independent pathways, suppressing IFNγ by lymphocytes and promoting IL-33 production by lung airway epithelial cells. However, a likely consequence of incoming L3 larvae is IL-33 release by damaged epithelial cells, which raises the question as to what are the circumstances that require IL-17A to further enhance IL-33 expression. A better understanding of the connection between IL-33 and IFNγ in this setting is therefore needed.

**Figure 2:**
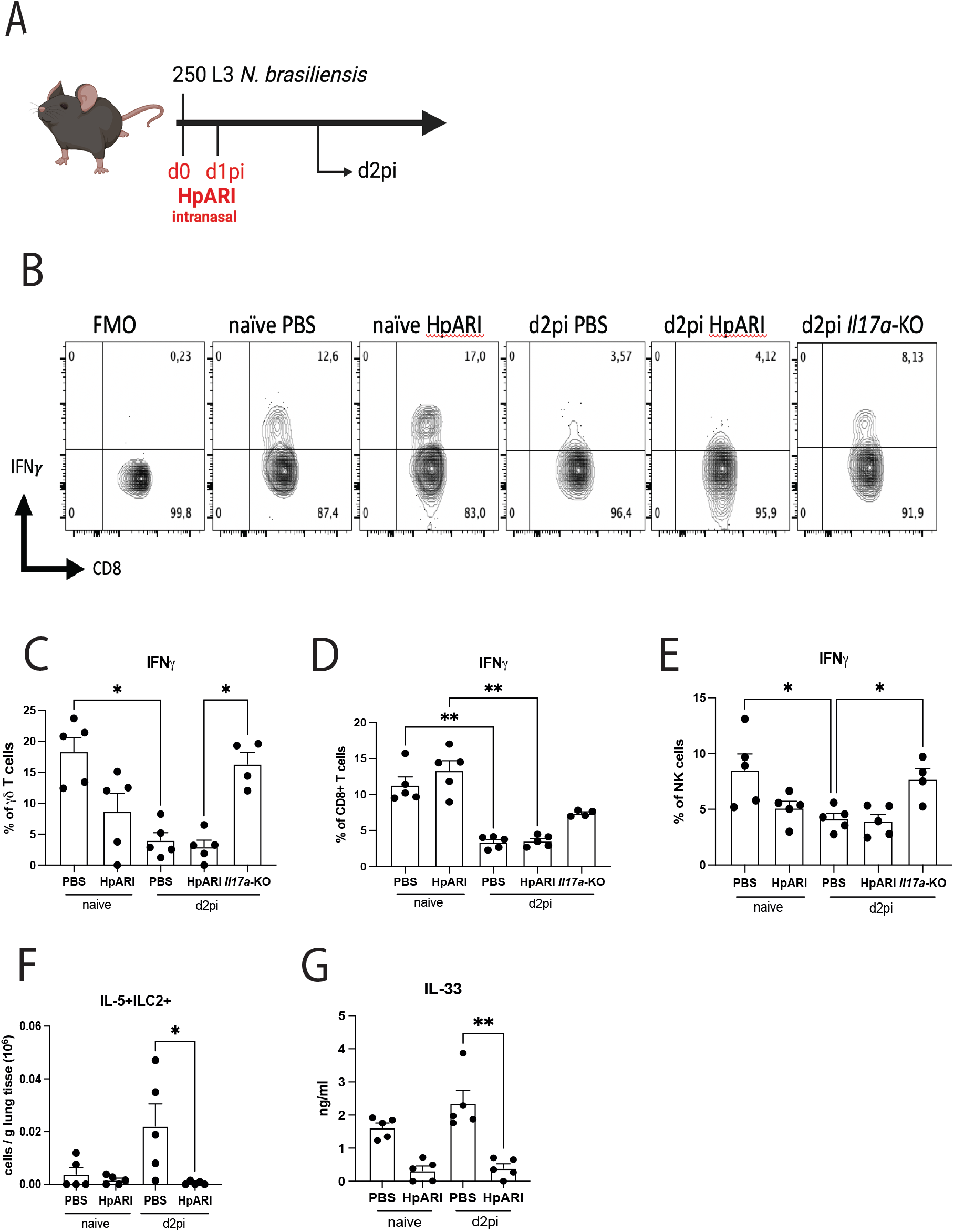
Mice were infected with 250 L3 larvae *Nb* and either treated with intranasal HpARI or PBS (A). Cellular sources of IFNγ were analysed via flow cytometry. Flow plots of IFNγ production by live TCRβ+CD8+ T cells in all tested groups as well as fluorescence minus one (FMO) control (B). Frequency of intracellular IFNγ by γδ T cells (C), CD8+ T cells (D), and NK cells (E) in naïve mice and mice infected with *Nb* d2pi, either treated with intranasal HpARI or PBS. *Il17a*-KO mice were used as positive controls. Absolute count of IL5+ILC2 in the lung in naïve and *Nb* infected mice (F). IL-33 in BAL fluid of naïve and mice d2pi as quantified via ELISA (G). Data are expressed as mean ± s.e.m. and are representative for 3 individual experiments and were analysed by ANOVA with Tukey’s multiple comparison test. \**P* < 0.05, \*\**P* < 0.01.

### IL-17A-dependent neutrophilia is not a driver of IL-33 production

A hallmark characteristic of IL-17A is its ability to promote neutrophil recruitment^8^. During the early phases of *Nb* infection, neutrophils are rapidly recruited to the lungs, with peak-neutrophilia being between d1 and d2pi^28^. Neutrophils primed by *Nb* infection have been shown to upregulate *Il33* transcripts^28^ and are a major driver of tissue injury which in turn could lead to IL-33 release. Therefore, we investigated whether neutrophils recruited during *Nb* infection contribute to IL-33 levels in the lung. Neutrophils were depleted at d-1 and 1pi (Fig. 3A) and successful depletion was confirmed via flow cytometry. Injection of anti-Ly6G effectively prevented neutrophil accumulation in the BAL at d2pi compared to isotype control (Fig. 3B). Histological sections of the lungs demonstrated infection-induced injury at d2pi in isotype-treated WT mice, an effect that is almost absent in infected mice depleted of neutrophils (Fig. 3C). Using measures of lacunarity as an indicator of acute lung injury^29^, neutrophil depleted mice displayed no significant signs of damage (Fig. 3C). These findings reinforce previous observations that neutrophilia is largely responsible for the severe lung damage observed in this model^28^. To investigate whether this extensive tissue damage during the early stage of *Nb* infection could be responsible for the increase in IL-33 release, lung sections were stained for IL-33 (Fig. 3D). Quantification of the IL-33 positive areas in the lung airways as well as the lung parenchyma did not reveal a difference between mice depleted of neutrophils and isotype controls (Fig. 3E, F). While infected mice exhibited increased IL-33 intensity compared to naïve mice, no significant difference was observed between the treatment groups. If neutrophils are a contributor of IL-33 during *Nb* infection, we would have expected less IL-33 after depleting the peak neutrophilia. However, mice depleted of neutrophils had a slightly higher intensity for epithelial IL-33 staining at d2pi, although this difference did not reach statistical significance (Fig. 3F). Depleting neutrophils also did not change the IL-33 levels at later stage of *Nb* infection as shown for d6pi in both lung epithelium and lung parenchyma (Fig. 3E, F). Our data therefore demonstrate that during the lung-migrating phase of *Nb* infection, neutrophil-mediated damage is not the cause of IL-33 production.

**Figure 3:**
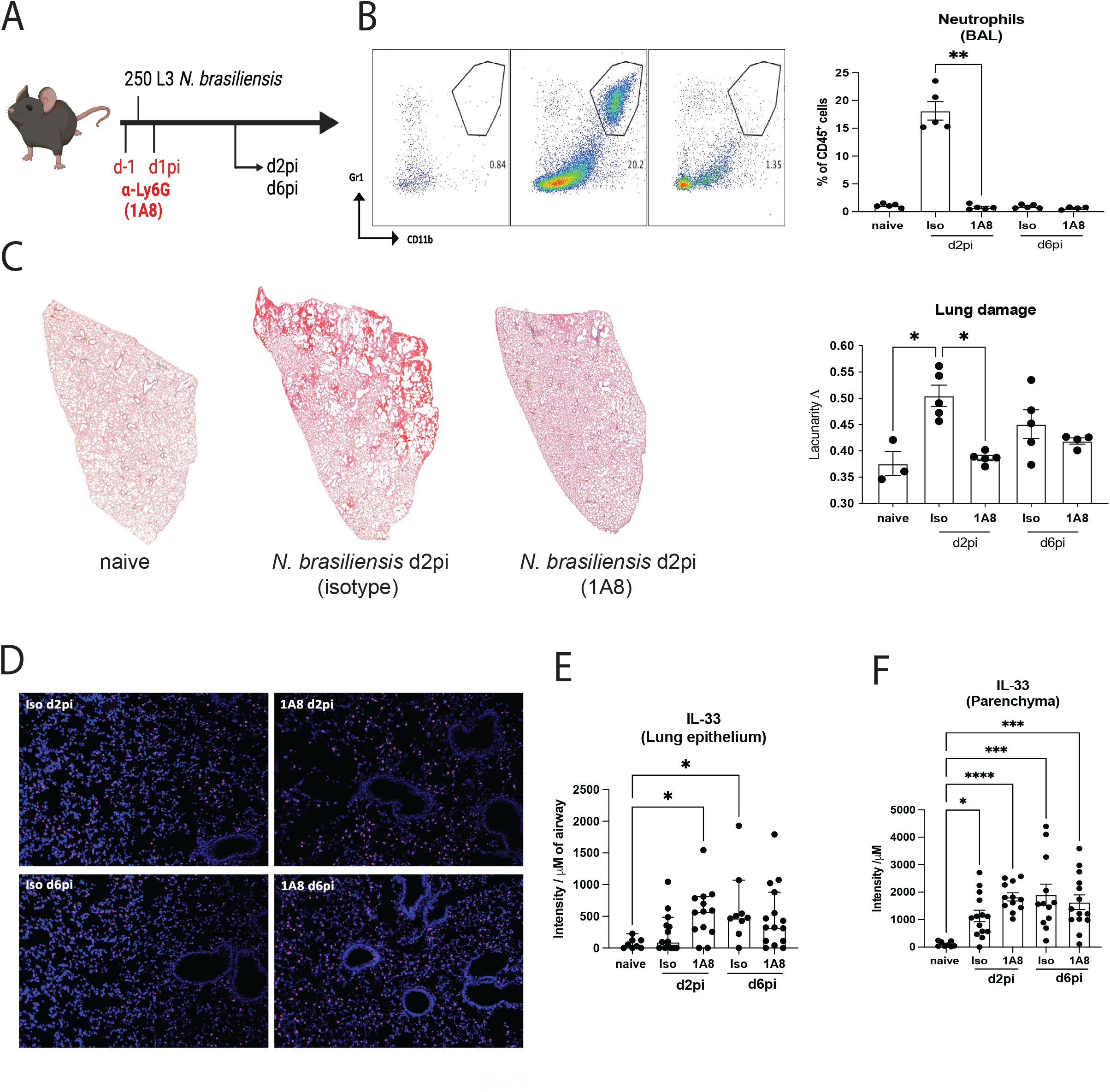
C57BL/6J mice were infected with 250 *Nb* L3 larvae and mice were injected intraperitoneally with α-Ly6G (1A8) or corresponding isotype control on d-1 and d1pi and immune responses were measured on d2pi or d6pi compared uninfected naïve mice (A). Confirmation of neutrophil (Gr1^+^ CD11b^+^) depletion via representative flow cytometry plots as well as neutrophil frequency in the BAL (B). Representative images of lung sections stained with hematoxylin & eosin and quantification of lacunarity (Λ) on d2pi and d6pi with *Nb* compared to lungs from uninfected naïve mice (C). Microscopy images of immunofluorescent staining for IL-33 (purple) in lung sections of mice depleted by neutrophils or isotype controls (D). Quantification for IL-33 positive areas for lung epithelium (E) and lung parenchyma (F). Antibody positive staining area was quantified for the intensity normalized to background staining. Data are expressed as mean ± s.e.m. and are pooled from 2 individual experiments and were analysed by ANOVA with Tukey’s multiple comparison test.

In summary, our brief report further implicates pulmonary IL-17A in the initiation phase of type 2 immunity in the lung. Through two independent mechanisms, enhanced IL-33 production by the lung epithelium, and suppression of IFNγ, IL-17A supports the type 2 immune response in the lung needed to cope with lung-migrating helminth infection.

## Acknowledgments

We thank the Flow Cytometry, Bioimaging, and Biological Services core facilities at the University of Manchester. This work was supported by the Wellcome Trust (106898/A/15/Z to JEA) with additional support from the Medical Research Council UK (MR/K01207X/1 to JEA) and the Wellcome Trust (221914/Z/20/Z to HJM)

## Author Contribution

J.A. executed experiments, S.P., J.E.P., and B.H.K.C. provided experimental assistance. J.A., T.E.S, J.E.A. designed experiments and analysed data, H.J.M. provided reagents and experimental advice. J.A., T.E.S. and J.E.A. wrote the original draft of the manuscript and all co-authors reviewed and edited the manuscript.

